# Genetic diversity and connectivity of the Ostreid herpesvirus 1 populations in France: a first attempt to phylogeographic inference for a marine mollusc disease

**DOI:** 10.1101/2021.04.30.442107

**Authors:** Jean Delmotte, Camille Pelletier, Benjamin Morga, Richard Galinier, Bruno Petton, Jean-Baptiste Lamy, Oliver Kaltz, Jean-Christophe Avarre, Maude Jacquot, Caroline Montagnani, Jean-Michel Escoubas

## Abstract

The genetic diversity of viral populations is a key driver of the spatial and temporal diffusion of viruses; yet, studying the diversity of whole genomes from natural populations still remains a challenge. Phylodynamic approaches are commonly used for RNA viruses harboring small genomes, but have only rarely been applied to DNA viruses with larger genomes. Here, we used the Pacific Oyster Mortality Syndrome (POMS, a disease that affects oyster farms around the world) as a model to study the genetic diversity of its causative agent, the Ostreid herpesvirus 1 (OsHV-1) in the three main French oyster-farming areas. Using ultra-deep sequencing on individual moribund oysters and an innovative combination of bioinformatics tools, we *de novo* assembled 21 OsHV-1 new genomes. Combining quantification of major and minor genetic variations, phylogenetic analysis and ancestral state reconstruction of discrete traits approaches; we assessed the connectivity of OsHV-1 viral populations between the three oyster-farming areas. Our results suggest that the Marennes-Oléron Bay represents the main source of OsHV-1 diversity, from where the virus has dispersed to other farming areas, a scenario consistent with current practices of oyster transfers in France. We demonstrate that phylodynamic approaches can be applied to aquatic DNA viruses to determine how epidemiological, immunological, and evolutionary processes act and potentially interact to shape their diversity patterns.

**Importance:** Phylogeography is a field of research that attempts to reconstruct the relationships between individual genotypes within a species and then correlate these genealogical relationships with their geographic and temporal origin. This field of research has become an essential step in the understanding of pandemics, in particular to determine the origin, spread and evolution of a pathogen as currently illustrated in studies on viral pandemics. However, because phylogeographic analyses are based on genome variation, stable genomes yield less information than labile genomes. Accordingly, viruses with double-stranded DNA (dsDNA) genomes generally have lower nucleotide diversity than RNA viruses. In this study, by combining the use of both major and minor genetic variations with phylogeographic analyses of the oyster herpesvirus OsHV-1, we highlight genealogical relationships that are not depicted in phylogenetic trees based on consensus viral genomes only. These data offer a plausible scenario reflecting the origin and spread of OsHV-1 populations between oyster-farming sites.

## Introduction

Viruses are disease agents that have high levels of genetic diversity. This high diversity often means that they lack shared genetic markers, such as ribosomal DNA sequences that are common to all prokaryotes and eukaryotes (Nkili-Meyong et al., 2016), making it difficult to characterize viruses genetically. Furthermore, many viruses - especially those with RNA genomes - are not stable genetic entities, but exist as clouds of many phylogenetically related genetic variations, known as viral quasispecies. This genetic organization currently encumbers our understanding of viral diseases and their evolution and it impedes the straightforward characterization of virus population structure (Lauring and Andino, 2010; Vignuzzi et al., 2006). Studies have shown that the level of genetic diversity within these viral populations likely influences viral pathogenicity, dissemination and host immune evasion. Full-length genome analyses are required to identify intra-host viral population structure, reveal molecular traits with epidemiological significance, or detect low-frequency, but nonetheless relevant viral variants. These difficulties apply to RNA viruses, but also to certain large DNA viruses, such as herpesviruses, whose genomic variability rivals that of many RNA viruses (Hammoumi et al., 2016; Renner and Szpara, 2018; Renzette et al., 2013). For example, human cytomegalovirus shows considerable inter-host and intra-host genetic divergence across tissue compartments and times of infection (Renzette et al., 2014; Renzette et al., 2015). In addition, the evolution of a disease is sometimes not fully explained by intrinsic host factors, but by the genotypic diversity of herpesviruses (Akhtar et al., 2019).

Aquaculture is one of the fastest growing food-producing sectors, representing an important animal protein supply for human consumption, with an expanding role in global food security. Today, the biggest threat arising due to the intensification and globalization of aquaculture is infectious diseases. The management and mitigation of the emergence and spread of these infectious diseases are key issues to address to ensure the sustainability of this industry (Burge et al., 2017; Pernet et al., 2016; Stentiford et al., 2012). One illustration is the Pacific oyster mortality syndrome (POMS), which threatens global *Crassostrea gigas* oyster production, a main sector in aquaculture worldwide (reviewed in (Petton et al., 2021)). Since 2008, this syndrome has caused mass mortality in cultivated oysters around the world, from Europe to America and Asia (Abbadi et al., 2018; Barbieri et al., 2019; Barbosa Solomieu et al., 2015; Friedman et al., 2005; Gittenberger et al., 2016; Jenkins et al., 2013; Moss et al., 2007; Peeler et al., 2012; Vasquez-Yeomans et al., 2010). In 2010, the causative agent of these massive mortalities was identified: it is an emerging genotype of a herpes-like virus named Ostreid herpesvirus 1 (OsHV-1) (Paul-Pont et al., 2013; Pernet et al., 2012; Petton et al., 2013; Renault et al., 2012; Segarra et al., 2010). Two major genetic factors seem to affect the severity of POMS: the genetic background of the oysters (de Lorgeril et al., 2018; de Lorgeril et al., 2020; Dégremont et al., 2015) and OsHV-1 genetic diversity (Delmotte et al., 2020; Friedman et al., 2020; Martenot et al., 2011). Since the characterization of the emergent genotype OsHV-1 µVar, associated with the 2008 high mortality events (Segarra et al., 2010), several variants of the µVar genotype have been described (reviewed in (Barbosa Solomieu et al., 2015)). Interestingly, in 2016, a survey of OsHV-1 genetic diversity carried out on wild *C. gigas* populations along the coasts of Italy demonstrated the high diversity of this virus in natural oyster populations (Burioli et al., 2016). However, most studies on OsHV-1 genetic diversity have been based on PCR molecular markers focusing on a few variable regions of the viral genome, which may not reflect the whole genomic diversity. For instance, in 2017, sequencing of the whole genome of OsHV-1 µVar and its comparison with the reference genome published in 2005 (Davison et al., 2005) showed that the two genomes also differed by the loss or addition of several open reading frames (ORFs), indicating that whole-genome sequencing is necessary to fully reveal viral diversity and to better understand the virus’ origin and evolution (Burioli et al., 2017). To date, no study has specifically looked at the links between the geographic distribution of OsHV-1 and its genetic diversity at whole genome level. As far as we know, the entire assembled genomes of only four OsHV-1 infecting *C. gigas* oysters are available, namely the reference genome (AY509253), two OsHV-1 μVar (µVar A KY242785.1 and µVar B KY271630.1) and more recently OsHV-1- PT (MG561751.2) (Abbadi et al., 2018; Burioli et al., 2017; Davison et al., 2005). These four assemblages show significant genomic diversity; however, they tell us very little about the geographic distribution and dissemination of the virus.

As shown, for instance, with the H1N1 influenza virus (Rambaut et al., 2008), the dengue virus (Allicock et al., 2012), the Zika virus (Theze et al., 2018), the AIDS virus (Faria et al., 2014) and more recently SARS-CoV-2 (Martin et al., 2021), molecular epidemiology based on full genomes can be the key to understanding the epidemics of emerging viruses. Therefore, the purpose of the present study was to characterize the genomic diversity in viral populations of OsHV-1 µVar encountered in oyster farming areas both within and between host individuals, to better understand the dissemination of OsHV-1 µVar populations and to assess the feasibility of applying phylodynamic and phylogeographic models to OsHV-1 whole genome sequences. Using a deep-sequencing approach conducted at the individual level and novel bioinformatics analyses; we identified 21 new complete genome sequences of OsHV-1 µVar from the three most important oyster-farming areas in France. Phylogenetic analyses combined with comparative genomics and inspection of variant frequencies showed that viral genetic diversity differs greatly not only between areas, but also between individuals within areas. An ancestral state reconstruction of discrete traits approach allows us to propose a scenario explaining the phylogenetic relationships between viral populations encountered in the three farming areas. This novel combination of approaches, together with epidemiological information, holds promise for understanding OsHV-1 evolution and epidemic dynamics and for identifying transmission patterns between oyster-producing regions. These data can be highly useful for developing novel disease management strategies.

## Results

### 1) Variability in sequencing depth does not impair the analysis of OsHV-1 diversity

We used specific-pathogen-free (SPF) *C. gigas* juveniles to sample OsHV-1 diversity during natural infections in the three farming areas (Fig. 1A). The cumulative mortalities for the three batches of oysters were 85%, 81% and 70% for Thau (Th), Marennes-Oléron (MO) and Brest (Br), respectively. No mortality was observed in the control group.

**Fig. 1:**
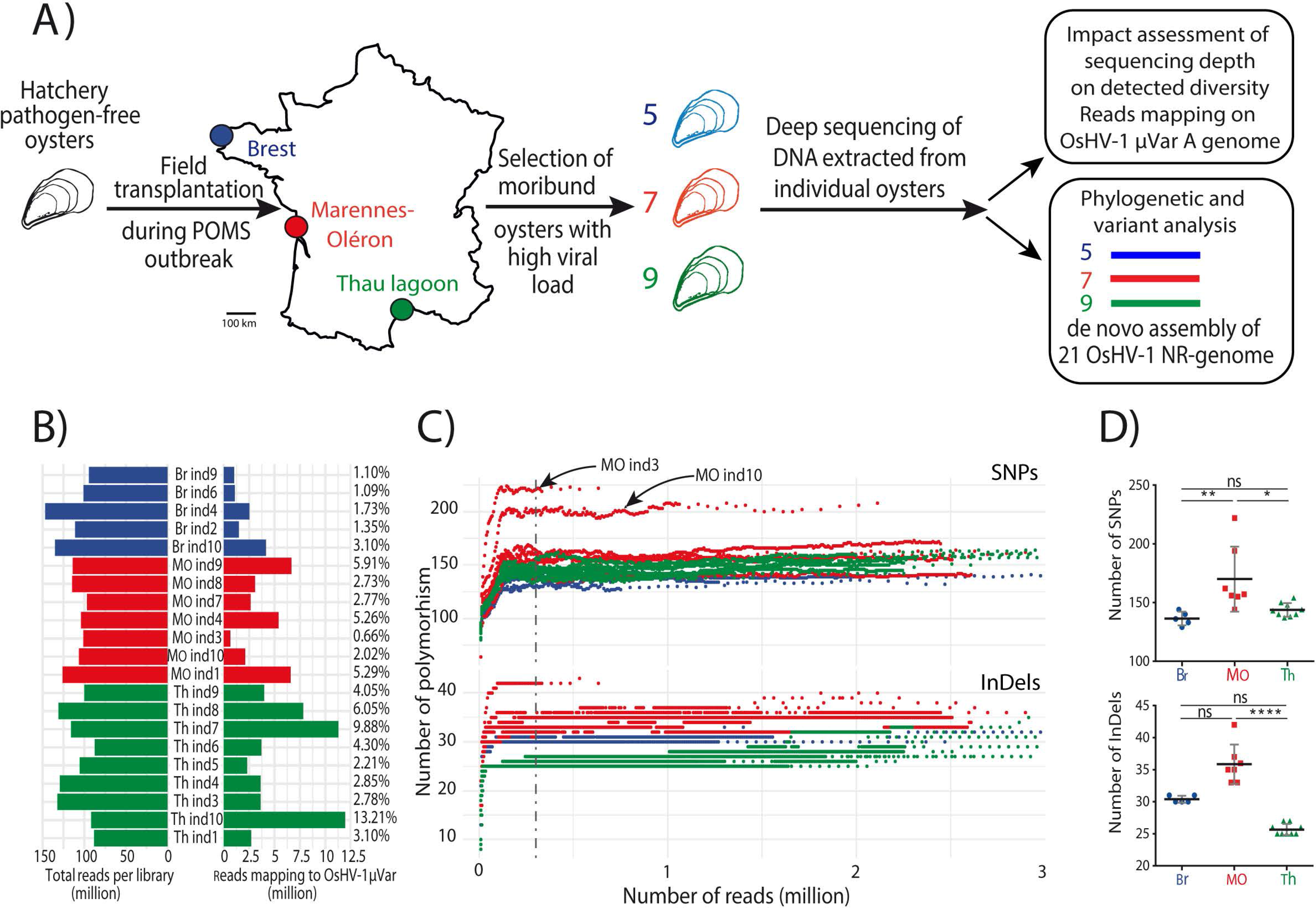
Variability in sequencing depth does not impair the analysis of OsHV-1 diversity. Schematic representation of the sampling of OsHV-1 genetic diversity in the three main French oyster farming areas during the Pacific oyster mortality syndrome (POMS) event in 2018 (See Material and Method for more details). Each color represents a farming area, with Brest (Br) in blue (5 individual samples), Marennes-Oléron (MO) in red (7 individual samples) and Thau Lagoon (Th) in green (9 individual samples). (**B**) Quantification of the number of reads in the library and the number of reads mapping to the OsHV-1 µVar A genome. Numbers on the right side of the figure indicate the percentages of reads mapping to the OsHV-1 µVar A genome. (**C**) Rarefaction curves showing the accumulation of polymorphisms (SNPs and InDels top and bottom panel respectively) during iterative variant calling analysis of the 21 libraries on the OsHV-1 µVar A genome. The dashed vertical line indicates 0.3 M reads, corresponding to the threshold at which the rarefaction curves reach a plateau. (**D**) Comparison of the number of SNPs (top) and InDels (bottom) between the three farming areas. Asterisks indicate the *P* value according to Kruskal-Wallis and Dunn’s multiple comparison tests (ns: non-significant, ∗*P* ≤ 0.05, ∗∗*P* ≤ 0.01 ∗∗∗∗ *P* ≤ 0.0001).

For each individual oyster, deep sequencing produced from 86.2 to 145.9 M reads per sample (average 109.7 M reads ±16.7 SD, Fig.1B. and Table S1). To evaluate OsHV-1 sequencing depth in each sample, sequencing reads were mapped on the OsHV-1 µVar A genome (KY242785.1) (Burioli et al., 2017). The number of reads mapping to the OsHV-1 µVar A genome greatly varied from one individual to the other, independently of the number of reads obtained for each sample: from 0.65 M reads (0.66%) for MO ind3 to 11.94 M reads (13.21%) for Th ind10 (Fig.1B, Table S1). Therefore, OsHV-1 genome coverage varied from 480X to 8745X for MO ind3 and Th ind10, respectively.

A rarefaction analysis was run using OsHV-1 µVar A as reference genome to verify that OsHV-1 coverage was sufficient to accurately quantify viral genomic diversity in each sample. For single nucleotide polymorphisms (SNPs) and for insertions-deletions (InDels), rarefaction curves reached a plateau around 300,000 reads, corresponding to an OsHV-1 coverage of 220 X (Fig. 1C). Since the number of mapped reads of each sample was clearly above this threshold, sequencing depth was considered sufficient to capture OsHV-1 genomic diversity. Rarefaction curves indicate that viral diversity was higher in infected MO oysters than in the other two areas. Comparison of the average number of individual polymorphisms deduced from rarefaction analysis between the three environments showed that i) the number of SNPs in viruses from MO (170 ± 27.7) was significantly higher than that of viruses from Br (136.4 ± 5.9; *P* ≤ 0.001) and Th (143.8 ± 5.8; *P* ≤ 0.05) and ii) the number of InDels in viruses from MO (35.9 ±3.1 SD) was also significantly higher than that of viruses from Th (25.7 ± 0.9; *P* ≤ 0.0001) (Fig. 1D, Table S2). In addition, in MO, viral diversity greatly varied from one individual to the other; for example, MO ind3 contained 222 SNPs and 42 InDels, whereas MO ind4 contained only 144 SNPs and 33 InDels.

### 2) Phylogenetic analysis of non-redundant OsHV-1 genomes reveals that all three OsHV-1 populations belong to the µVar genotype

We assembled non-redundant (NR) genomes so as to reduce the complexity of the OsHV-1 genome and keep only one copy of each repeated region in the virus sequence. The OsHV-1 µVar A genome contains two unique regions and three repeats: the unique long region (UL, 164,268 base pair (bp)), flanked by two inverted repeats (repeat long [RL], 7338 bp), and the unique short region (US, 3370 bp), flanked by two inverted repeats (repeat short [RS], 9777 bp), and two X regions (1510 bp) (Fig 2A, upper panel). Considering the high sequence identity within each set of repeats (between 99.9 and 100%), removing one of the repeats makes it possible to reliably map the reads from these repeated regions without affecting the overall genome composition. Removing a copy of each repeated region led to a NR OsHV-1 µVar A genome (NR-genome) of 186,262 bp, which represents 91% of the full-length genome and contains all the coding sequences (Fig 2A, lower panel).

**Fig. 2:**
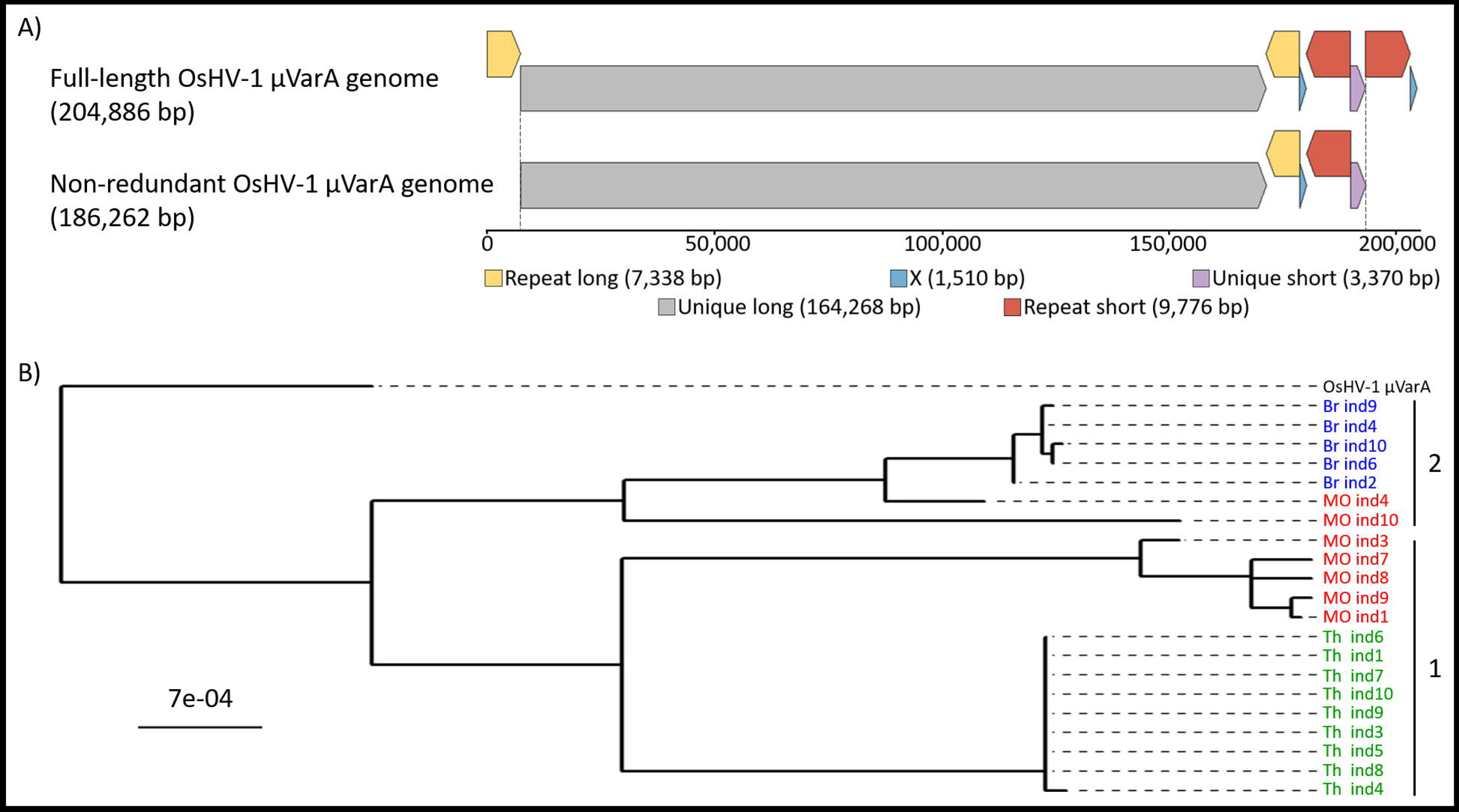
Phylogenetic relationship between the 21 *de novo* assembled and other OsHV-1 non-redundant genomes. (**A**) Full-length (upper panel) and non-redundant (NR) (lower panel) genomes of OsHV-1 µVar A (KY242785). The OsHV-1 µVar A genome consists of two unique regions, referred to as unique long (UL; in grey) and unique short (US, in purple), and three repeat regions, referred to as repeat long (RL, in yellow), repeat short (RS, in orange) and X region (X, in blue). The first two repeat regions each have copies at a terminal locus (TRL, TRS) and an internal locus (IRL, IRS) within the genome. The NR-genome consists of the linear fragment starting at the beginning of the UL and ending at the end of US region. This strategy includes all the genetic information without duplication. All the regions within the genome and the NR genome are to scale. (**B**) Shown is a maximum-likelihood phylogenetic tree of the 22 OsHV-1 NR-genomes rooted in OsHV-1 µVar A. The 21 isolates from this study are colored according to their geographic origin, Brest (Br) in blue, Marennes-Oléron (MO) in red, and Thau Lagoon (Th) in green. Within those, two clusters were defined based on genetic distances (numbered on the right).

Using this approach, 21 NR-genomes were assembled *de novo* from the sequencing data generated during this study, as well as 6 other *Ostreid herpesvirus 1* (belonging to the *Malacoherpesviridae* family and *Ostreavirus* genus) published genomes. A phylogenetic analysis of these 27 NR-genomes revealed that the 21 new OsHV-1 isolates were phylogenetically closer to OsHV-1 µVar A and B isolates (Fig. S1) than to the OsHV-1 reference genome (AY509253.2), suggesting that they belong to the µVar genotype.

Among newly sequenced isolates, those originating from Th and Br grouped into two distinct clusters reflecting their geographic origin (Fig. 2B, clusters 1 and 2). Conversely, MO OsHV-1 isolates were distributed over these two clusters: five in cluster 1 and two in cluster 2. The phylogenetic distance among Th or Br isolates (average 2.47 10^-5^ ± 4.68 10^-5^ SD and 1.43 10^-4^ ± 7.91 10^-5^ SD, respectively) was lower than that observed among the MO isolates (average 5.00 10^-3^ ± 4.21 10^-3^), indicating greater viral diversity in MO than in the other two farming areas.

### 3) Comparative genomics of the 21 non-redundant genomes confirms that OsHV-1 diversity is higher in Marennes-Oléron than in the other two farming areas

To compare the genetic diversity at the whole-genome scale, we used the 21 NR-genomes to construct an average non-redundant genome (A-genome) (See Materials and Methods for more details). The comparison of the 21 NR-genomes to the A-genome identified a total of 399 genetic variations (SNPs and InDels, hereafter called variations), spread over 310 positions. Twenty two of these positions contained from 2 to 7 allelic variations (Table S3). Prediction of the possible functional effects of the 399 variations revealed that 216 (54.1%) were distributed over 94 ORFs (summarized in Fig. S2 and detailed in table S3). The 21 variations predicted to have a high impact (frameshift or start lost) were distributed over 11 ORFs. Unfortunately, these results are not very informative given that 6 ORFs were annotated “unknown” due to the absence of homologies with any known proteins and the 5 others were annotated according to domain homologies suggesting a protein localization or a putative function.

The average number of variations per genome was significantly lower in samples from Th (63.7 ± 2.7) than from MO (92.1 ± 25.8; *P* < 0.05) and Br (120.8 ± 2.8; *P* < 0.001), whereas the average number of variations in MO was not significantly different from Br (Fig. 3A, Table S3). However, viruses from MO showed the highest diversity, with 290 variations accounting for 72.7% of the total diversity. Among the variations, 187 (64.5%) were specific to MO, and 114 were observed in one sample only (singletons) (Fig. 3B, Table S3). None of these 187 variations were shared by all the samples from this area. The majority of MO singleton variations were observed in two samples: ind10 (56) and ind4 (23) (Fig. 3C). For Th and Br farming areas, 82 and 132 variations were identified, among which 70 and 38 were area-specific, respectively (Fig. 3B, Table S3). In contrast within MO, 57.9% (22/38) and 75.5% (53/70) of the variations were common to all the samples of Br and Th, respectively (Fig. 3B, Table S3). Among the 92 variations shared by Br and MO, 87 (94.6%) were shared by two MO individuals (ind10 and ind4) and all Br individuals (Fig. 3C, purple arrow). These two MO individuals also shared five variations with all Th individuals (Fig. 3C, red arrows).

**Fig. 3:**
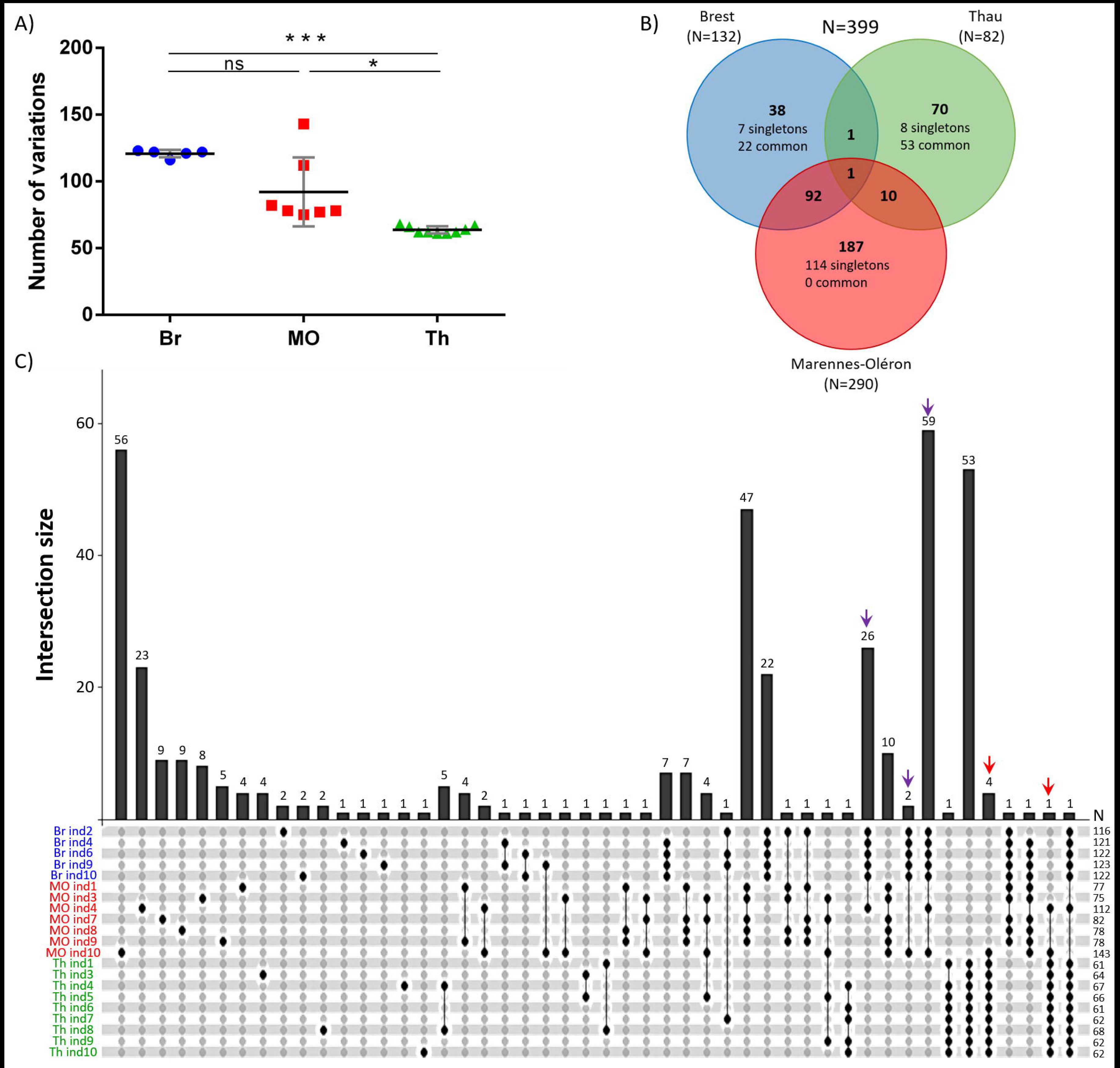
Comparative genomics of the 21 non-redundant genomes of OsHV-1 isolated from three oyster farming areas. (**A**) Distribution of the 399 variations between the 21 individuals and the three farming areas. Significance was calculated according to Kruskal-Wallis and Dunn’s multiple comparison tests (ns: non-significant, ∗ *P* < 0.05, ∗∗∗ *P* < 0.001). (A) Venn diagram summarizing the distribution of variations between the three farming areas. Color depends on the farming area with Brest (Br) in blue, Marennes-Oléron (MO) in red and Thau Lagoon (Th) in green. The numbers of variations per farming area are indicated in parentheses. Singletons are variations that occur in only one sample, and the term “common” indicates variations present in all the sampled individuals at a given location. (**C**) UpSet plot of variation distribution among the 21 non-redundant (NR-) genomes. The bottom panel shows the specific combinations (or intersections), and the vertical bars indicate the number of variations within these combinations. Colored arrows on top indicate variations shared by two individuals from MO (ind4 and ind10) and all individuals from Br (purple arrows) or all individuals from Th (red arrows). Numbers (N) on the right side indicate the number of variations per sample.

Interestingly, these major variations were only characterized in a small number of individuals within the samples analyzed; indeed, 129 (32%) of them are found in only one individual and only 2 (0.5%) are shared by more than 50% of individuals.

### 4) Within-individuals diversity of OsHV-1: the minor variations are more frequent in Marennes-Oléron than in the other two farming areas

To fully characterize OsHV-1 genetic diversity, we performed variant calling analyses, using the A-genome as the reference, to quantify both major (frequency >50%) and minor (frequency <50%) variations within each of the 21 individual samples. Among the 399 variations identified as major variations, 132 had variable frequencies (above or below 50%) across samples; moreover, 208 additional minor variations were identified (Fig. 4A, Table S4). These 208 minor variations are spread over 164 positions of the A-genome and 17 of these positions are showing 2 to 12 allelic variations (Table S4). SnpEff predicts that 72 (18.0%) of the 208 variations were distributed over 34 predicted ORFs (summarized in Fig. S3 and detailed in table S4). The 7 variations predicted to have a high impact (frameshift or start gained) were distributed in 6 ORFs. But, again, these results were not very informative given that 4 of these ORFs were annotated “unknown” and the 2 others were annotated according to domain homologies suggesting a membrane location of the predicted protein.

**Fig. 4:**
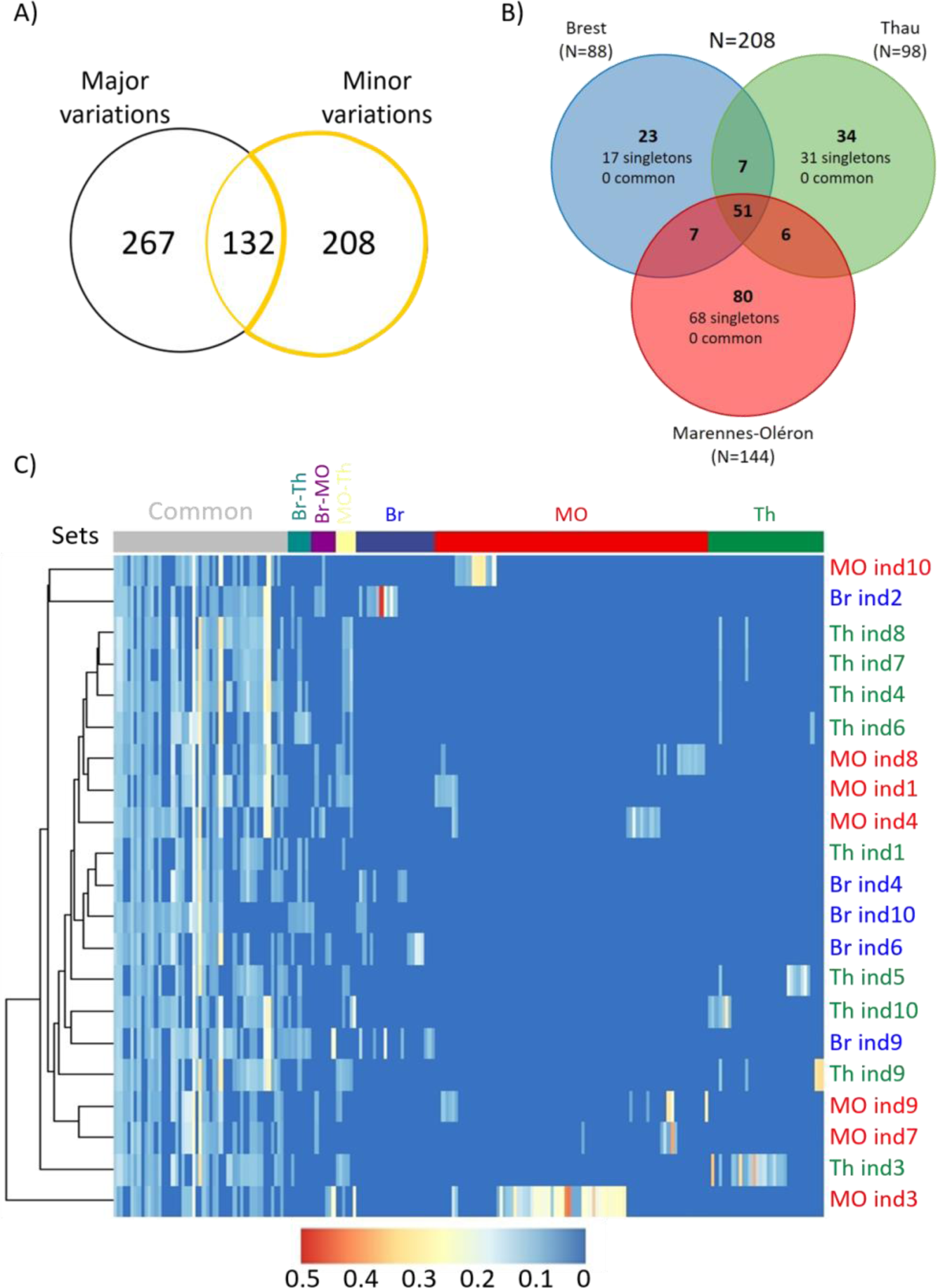
Variant calling analysis and minor variation distribution in the three farming areas. (**A**) Quantification of both major and minor variations using a variant calling analysis using A-genome as the reference (**B**) Venn diagram summarizing the distribution of the 208 strictly minor variations (frequency <50%) between the three farming areas (Brest (Br) in blue, Marennes-Oléron (MO) in red, and Thau Lagoon (Th) in green). The number of minor variations within each farming area is indicated in parentheses. Singletons correspond to variations that occur in only one oyster sample, whereas the term “common” corresponds to variations present in all the oyster samples within the dataset. (**C**) Heatmap of the allelic frequency of strictly minor variations across the 21 samples with a gradient from 0 (blue) to 0.5 (red). Hierarchical clustering using the Euclidean distance of the allele frequency is displayed on top. Each line of the heatmap corresponds to a strictly minor variation. variation sets on the top are derived from the Venn diagram. Samples are indicated on the right, and colored according to their geographic origin.

Frequencies of the 208 strictly minor variations ranged from 5.0% to 48.7% (mean = 9.57% ± 6.52% SD). The number of minor variations was lower in oysters from Br (88) than in oysters from Th (98) or MO (114) (Fig. 4B, Table S4). Fifty-five of these 208 minor variations (24.5%) were found in all three farming areas. Moreover, all site-specific variations were found in only one individual oyster (singletons).

A hierarchical clustering analysis applied to the minor variations showed that individuals did not cluster according to their geographic origin (Fig. 4C). In some cases, we even observed two individuals from different farming area clustering together (MO ind10 and Br ind2, Fig. 4C), suggesting that some OsHV-1 found in these individuals had a common origin. Figure 4C also revealed that the 51 variations shared by the three farming areas (common) were found in a majority of sampled individuals (mean = 13.9 ± 5.7 SD), suggesting that they had not arisen within individual oysters, but had been transmitted horizontally to other oysters.

### 5) Distribution of within and between individual genetic variations among samples reveals oyster farming areas connectivity

The 132 variations that had variable frequencies across samples (Fig. 4A) were present as minor variations in only four samples: three from MO (ind3, ind4, and ind10) and one from Br (ind2) (Fig. 5A). Among them, samples MO ind3 and MO ind10 contained 125 of these minor variations. Sample MO ind10 contained 30 minor variations (with a frequency ranging from 5 to 7.2 %; Fig. 5B, Table S5) that were major variations in samples MO ind1/3/7/8/9. For instance, the variation at position 65,003 (T>A) of the A-genome had a frequency of 7% in MO ind10, whereas its frequency varied from 76% to 99% in MO ind1/3/7/8/9 (Fig. 5C). Similar fluctuations in variation frequency were observed for the other 29 variations (Table S5). These 30 variations highlighted a clear genetic connectivity between the MO ind10 sample and the five MO samples (ind1/3/7/8/9) located in the other phylogenetic clade (Fig. 5D).

**Fig. 5:**
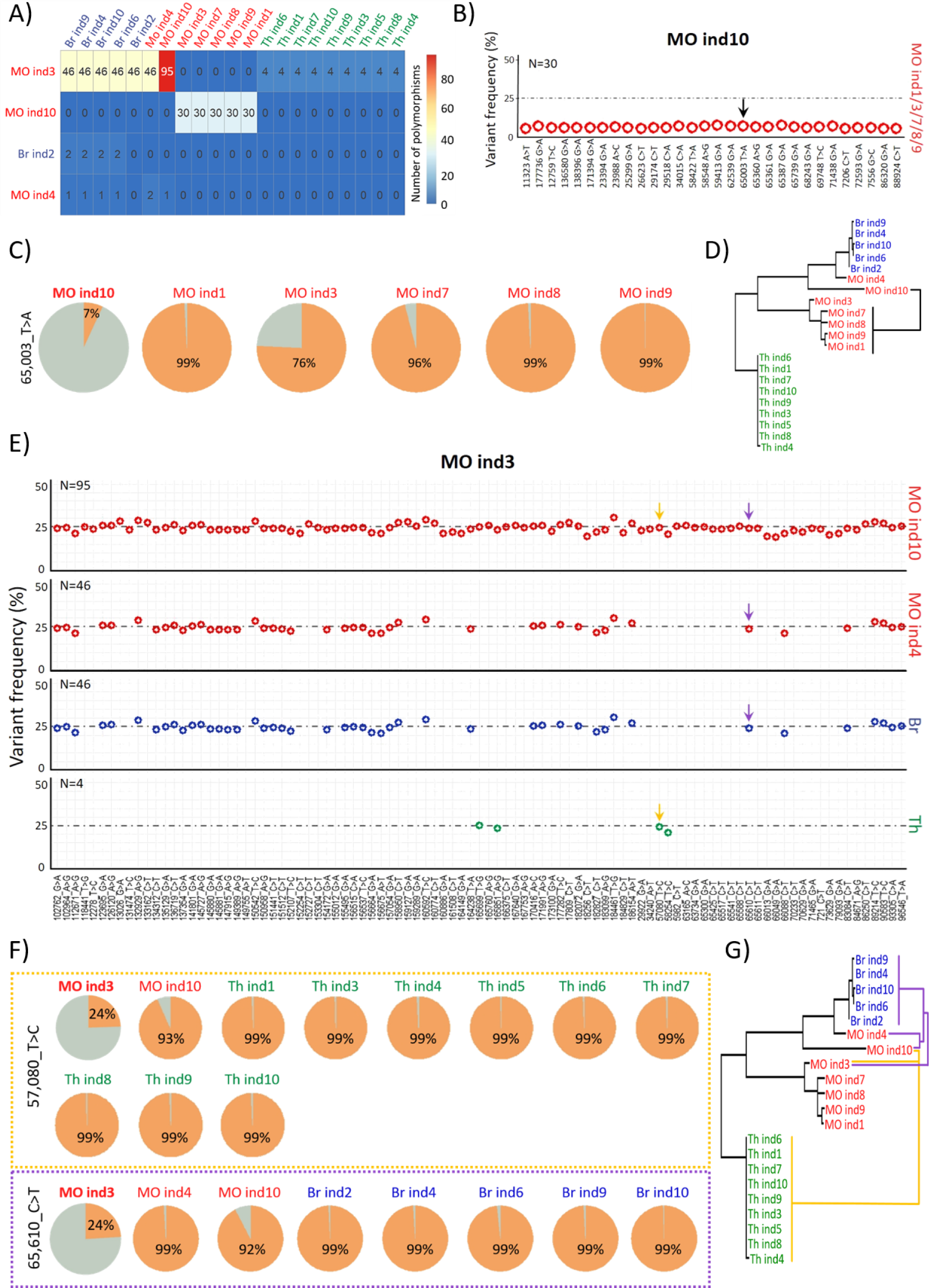
Some minor OsHV-1 variations found in the Marennes-Oléron farming area are major variations in the other two farming areas. (A) Matrix of the 132 variations which were major or minor, according to the sample. Rows display the four samples in which the 132 variations were minor, and columns show all 21 samples. Intersections give the number of shared variations with minor (row) and major (column) frequency. For self-comparisons, no number is given. (**B**) Dot plot of the 30 minor variations of MO ind10 which were major variations in MO ind1/3/7/8/9. Positions on the A-genome and the type of SNPs are indicated on the x-axis. variation frequency (in %) is indicated in ordinate. Black arrow indicates the variation that is detailed in (C). (**C**) Pie charts of the frequency of the variation found at position 65,003 of the NR-genome in MO ind1/3/7/8/9/10. (**D**) Phylogenetic tree of the 21 *de novo* assembled NR-genomes rooted in midpoint. The black line shows the genetic connectivity established between MO ind10 and MO ind1/3/7/8/9 from variation frequencies at the 30 positions indicated in panel B. (**E**) Dot plot showing the position and type of SNPs (x-axis) and variation frequency (y-axis) of the 95 minor variations of MO ind3 that were present as major variations in MO ind10 (95 variations), in MO ind4 (46 variations) as well as in all individuals from Br (46 variations) and Th (4 variations). Yellow and purple arrows indicate the two variations (57,080_T>C and 65,610_C>T respectively) that are detailed in (F). (**F**) The minor variation found in MO ind3 at position 57,080 of the A-genome corresponds to major variations in MO ind10 and all Th samples (upper panel, framed in yellow). The minor variation found in MO ind3 at position 65,610 of the A-genome corresponds to major variations in MO ind4/10 and all Br samples (lower panel, framed in purple). (**G**) Phylogenetic tree of the 21 *de novo* assembled NR-genomes rooted in midpoint. The yellow lines show the genetic connectivity established between MO ind3 and MO ind10 and all Th samples based on the study of variation frequencies at the 4 positions indicated in panel E. The purple lines show the genetic connectivity established between MO ind3 and MO ind4, MO ind10 and all Br samples based on the study of variation frequencies at the 95 (MO ind4) and 46 (MO ind10 and all Br samples) positions indicated in panel E.

Furthermore, MO ind3 contained 95 minor variations that corresponded to major variations in MO ind10, 46 that corresponded to major variations in MO ind4 and all Br samples, and 4 that corresponded to major variations in all Th samples (Fig. 5 A and E). For instance, the variation at position 57,080 (T>C) in the A-genome had a frequency of 24% in MO ind3, whereas its frequency was 93% in MO ind10 and over 99% in all Th samples (Fig. 5F, Table S5). Another example is the variation at position 65,610 (C>T) in the A-genome: its frequency grew from 24% in MO ind3 to 92% in MO ind10, and 99% in MO ind4 and allBr samples (Fig. 5F, Table S5). Again, minor variations highlighted genetic connectivity between samples from different farming areas and from phylogenetic clusters 1 and 2 (Fig. 5G, Table S5).

### 6) Ancestral state reconstruction of discrete trait

The analysis of minor variations highlighted links not only between MO samples and samples from the other two farming areas, but also between MO samples, which segregated into two different clusters according to whole-genome phylogenetic analyses. These results indicate that all the OsHV-1 variations characterized in this study originated from the MO farming area. This is congruent with ancestral state reconstruction of the discrete locations (Fig. 6A, Fig. S4) that showed evidence for MO being the ancestral location of the 21 OsHV- 1 isolates sequenced. Moreover, the Bayesian stochastic search variable selection approach suggested significant transition rates of OsHV-1 between the three sampling sites regardless of the alignment used to perform the analysis, *i.e.* full NR-genomes or NR-genomes with regions potentially impacted by recombination events removed (Table S6). Indeed, two rates were inferred with decisive supports: from MO to Th (BF=3683) and from MO to Br (BF=11,051) (Fig. 6B).

**Fig. 6:**
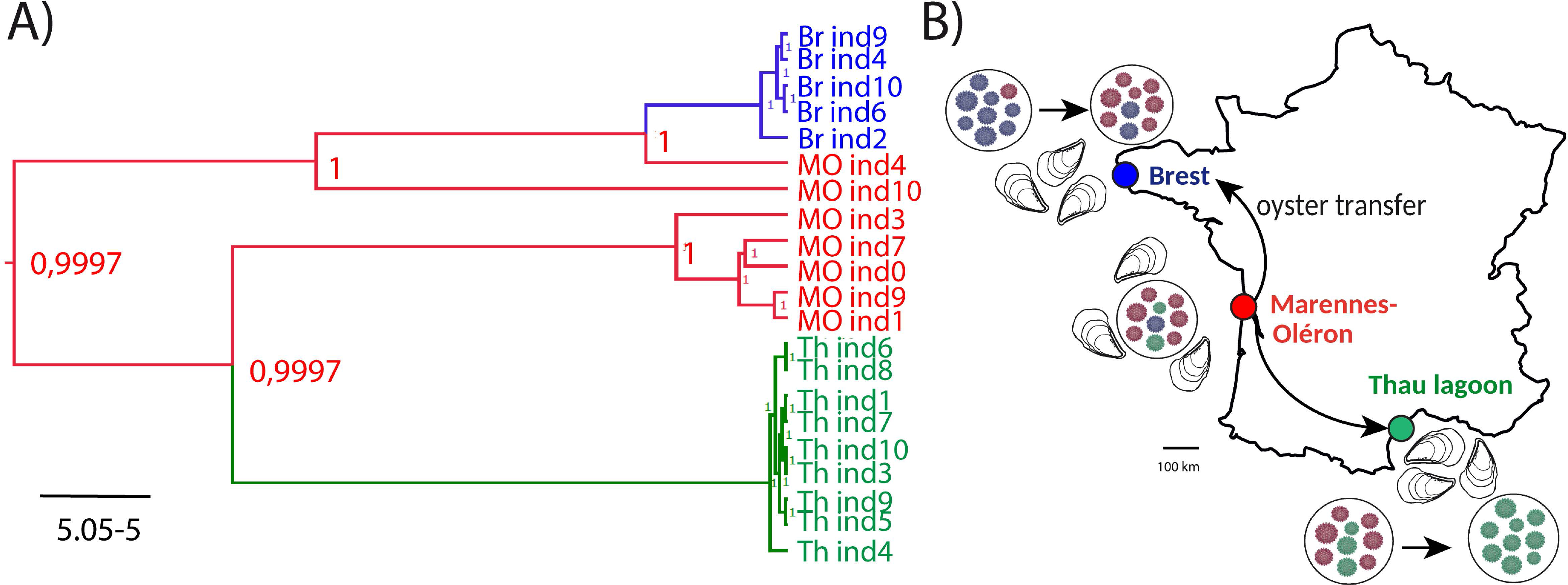
Graphical representation of OsHV-1 population connectivity in France. **(A)** Maximum clade credibility (MCC) phylogeny of NR-genomes of OsVH-1 isolates and ancestral state reconstruction of the discrete locations: Brest (Br), Marennes-Oléron (MO) and Thau lagoon (Th). Color of the branches represents the most probable location state of their descendent nodes. The color-coding is consistent with other figures: Br in blue, MO in red, and Th in green. Numbers at branch nodes indicate the posterior location probabilities. Given that we did not incorporate any timing information into our analysis (no calibration prior), the scale beneath the tree is expressed in average number of substitutions per time unit. (**B)** Bayesian stochastic search variable selection (BSSVS) was used to infer dispersal routes with none-zero rates using Bayes factors. Routes that were inferred to have decisive support (Bayes Factor > 100) are plotted on map (black arrows).

## Discussion

The accurate description of viral genetic diversity (major and minor variations) depends on several parameters, such as the quantity and quality of viral genetic material, sequencing quality and depth, as well as the bioinformatics tools used for data analyses (Sanjuán and Domingo-Calap, 2019). In the present study, we characterized OsHV-1 genetic diversity in three oyster-farming areas during POMS outbreaks. To do so, oysters from the same cohort were exposed to POMS outbreaks in the field and OsHV-1 genetic diversity was investigated in individual infected moribund oysters. To optimize the characterization of OsHV-1 variations, we selected moribund oysters with high viral loads. Then, DNA extracted from these oysters was used to prepare sequencing libraries using a PCR-free kit to remove genomic coverage biases associated with PCR amplification. Using this approach, we obtained an average OsHV-1 genome coverage of ≈3000X. However, because OsHV-1 coverage varied among samples (ranging from 480X to 8745X), it was necessary to find a way to verify that coverage was sufficient to accurately quantify viral genomic diversity in each sample. Using a rarefaction analysis, we determined the threshold at which sequencing depth was sufficient to capture all of the genetic diversity present in the studied samples. Although this analysis is commonly used in ecology to assess the species richness in a community (Chase et al., 2018), it is, to our knowledge, rarely applied to study viral genetic diversity. This information can be crucial when studying viral diseases in which the high diversity of the virus may potentially increase the fitness of the viral population, making it hard to eradicate (Al Khatib et al., 2020; Domingo et al., 2012).

Having a great sequencing depth was an advantage for OsHV-1 genome assembly; however, the task was complicated by the presence of large repeated sequences, corresponding to 9.1% of the genome of OsHV-1 µVar A. To solve this problem, we used an approach commonly used for herpesvirus genome analyses (Cunningham et al., 2010; Morse et al., 2017): contigs generated by *de novo* assembly were ordered using a reference-based approach to ultimately assemble non-redundant genomes in which only one copy of each repeated element was kept. These non-redundant genomes constituted the keystone of our bioinformatics analyses. Because each of them contained the entire genomic sequence of the virus, we were able to carry out an exhaustive search for intra-individual (sample) genetic variations and characterize the viral population structure. These genomes were also aligned with each other to perform whole-genome phylogeny and genome-wide comparative genomics.

Our results revealed that viral populations of OsHV-1 are a heterogeneous set of genomes rather than a single dominating genome, as revealed for another aquatic herpesvirus, the Cyprinid herpesvirus 3 (Hammoumi et al., 2016). They also suggest that all OsHV-1 characterized in our study share a common ancestor with OsHV-1 µVar characterized by Burioli *et al*. in Normandy (France), reinforcing the idea that genotypes closely related to µVar have replaced the reference genotype and are predominant in oyster-farming areas along the French coasts (Burioli et al., 2017; Burioli et al., 2018). Our findings also showed that OsHV-1 µVar populations are much more diverse than previously thought. This is congruent with results from Delmotte *et al*. showing that OsHV-1 µVar from the Bay of Brest and Thau Lagoon constitute two distinct viral populations (Delmotte et al., 2020) as well as with the recent relatively high estimate of OsHV-1 evolutionary rate (Morga et al., 2021). We therefore argue it is now necessary to combine common approaches relying on the generation and the analysis of consensus genomes to intra host diversity characterization in order to accurately study the epidemiology of DNA viruses (Renner and Szpara, 2018).

Here, we identified major and minor variations within viral populations tha substantially vary in frequency among sampled oyster individuals. Thus, ultra-deep sequencing can support the reconstruction of the true structure of a viral population, and inform on relationships between individuals from the same farming area or between individuals from the three farming areas (Fig. 5 D and G, respectively). Ancestral state reconstruction, used to infer the dynamics of dispersal of OsHV-1 populations across the three farming areas (Fig. 6 A), along with the distribution of major and minor variations, suggests that Marennes-Oléron farming area is the main source of OsHV-1 viral diversity, from where variations are dispersed to the other farming areas. This hypothesis is in agreement with what is known about oyster transfers. Indeed, Marennes-Oléron is the main area of *C. gigas* spat collection, and juvenile oysters are then transferred to growing sites along the coasts of Brittany, Normandy and also along the Mediterranean (Buestel et al., 2009) (Fig. 6 B). Importantly, the three studied sites are separated by long distances (>500 km) and therefore are not directly connected through ocean currents. OsHV-1 thus likely disseminates via the transportation of infected oysters, similar to the spread of other diseases via livestock transfer (*e.g.*, (Keeling et al., 2001)). Effectively several studies showed that wild-caught spats, or oysters surviving an OsHV-1 infection contain the virus and are therefore able to transmit it when cohabiting with uninfected oysters (Agnew et al., 2020; Dégremont and Benabdelmouna, 2014; Schikorski et al., 2011).

OsHV-1 diversity found in Marennes-Oléron Bay was higher than in the Bay of Brest and Thau Lagoon. This may be related to the fact that the Marennes-Oléron Bay has become one of the largest French oyster-farming areas, with intensive shellfish cultivation and chronic overstocking (Buestel et al., 2009). While we acknowledge the potential bias resulting from small sampling size, this is coherent with large viral population size and increased opportunities for recombination. Mutation and recombination rates being good prognostic markers for possible emergence of phenotypically different viruses, studying them at the different farming areas would help to better prevent future new OsHV-1 variation outbreaks. Moreover, a network analysis of oyster movements revealed that oysters are moved up to 9 times during their production cycle, with peaks of transfers in spring and autumn. These aquacultural practices likely contribute to the spread of OsHV-1, and in turn increase its genetic diversity (Lupo et al., 2016).

Given that the fate of new genetic variations is largely determined by host selection, one hypothesis to explain the lower diversity in the Bay of Brest and Thau Lagoon is that oyster populations there exert a bottleneck on viral diversity imported from Marennes-Oléron. Indeed, it can be speculated that some OsHV-1 variations imported from Marennes-Oléron have not found oysters with a genetic background compatible with their replication and survival. However, several studies conducted in Europe to document oyster genetic variability and population structure have revealed a high level of genetic diversity, but no genetic differentiation between French populations (Lapegue et al., 2020; Rohfritsch et al., 2013; Vendrami et al., 2019). This lack of differentiation suggests that selection pressure exerted by oysters is not sufficient to explain differences in viral genetic diversity between farming areas. Genetic drift also influences the probability and rate by which alleles increase or decrease in frequency in a viral population; drift results in the loss of genetic diversity, because only a subset of the population contributes to the next generation (Sanjuán and Domingo-Calap, 2019). Furthermore, the relationship between environmental factors and mortality events in Pacific oysters has been well documented (reviewed in (Alfaro et al., 2018)). Environmental factors such as temperature, food availability, water quality and salinity or UV radiation have been identified as risk factors that directly or indirectly affect POMS dynamics. Given the variation in these factors among the studied oyster farming areas, identifying the effects of these factors on the structure of OsHV-1 µVar populations may shed new light on the processes of variation selection. Therefore, a larger OsHV-1 study should focus on characterizing the epidemiological and evolutionary processes that shape OsHV-1 genetic diversity patterns and on identifying the ecological, environmental and anthropogenic factors influencing OsHV-1 dispersal using a combination of phylogeography and spatial epidemiology approaches (Dellicour et al., 2016; Pybus et al., 2012).

Combining data on OsHV-1 viral population structure using an innovative combination of bioinformatics tools with robust epidemiological information from the different farming areas can offer valuable insight into the dynamics of POMS infection (routes of transmission, potential reservoirs) and thus ultimately help implement effective integrated viral disease management strategies (Avarre, 2017).

## Materials and Methods

### OsHV-1 field infection and oyster sampling

All *Crassostrea gigas* oysters used in the present study were produced in August 2017 in the Ifremer experimental facilities located in Argenton (Brittany, France), as specific-pathogen-free (SPF) juveniles (Petton et al., 2015). This SPF oysters cohort was generated from 132 genitors (108 females and 24 males) derived from wild-caught spat collected between 2012 and 2015 (33 genitors for each year). To sample OsHV-1 diversity, SPF oysters originating from the same batch were transplanted into three oyster farming areas (∼1000 individuals per farm; average weight 1.5 g/oyster) during a disease outbreak when seawater temperature was above 16°C: 1) the Thau Lagoon (Th) (Mèze, lat. 43.379 long. 3.571) in May 2018, 2) the Marennes-Oléron Bay (MO) (La Floride, lat.: 45.803 and long.: - 1.153) in June 2018 and 3) the Bay of Brest (Br) (Logonna-Daoulas, lat.: 48.335 long.: -4.318) in July 2018 (Fig. 1A). It has been established that massive mortalities can occur as early as 5 days after oyster transplantation in the natural environment (Dupont et al., 2020; Petton et al., 2015). For this reason, oysters were transferred back to laboratory facilities 5 days after transplantation. They were then maintained in seawater tanks at 20°C (density of 20 to 25 oysters/L), and moribund oysters were collected daily and stored at -80°C until analysis. A control group was composed of oysters that had not been transplanted in natural environment. Monitoring was stopped after three consecutive days without any oyster mortality. During transport, oysters were packed in polystyrene containers and kept moist by covering them with a damp cloth.

### DNA extraction, viral load quantification and sequencing

DNA was extracted from individual moribund oysters using the MagAttract^®^ HMW DNA kit (Qiagen) according to the manufacturer’s protocol. DNA purity and concentration were checked using a Nano-Drop ND-1000 spectrometer (Thermo Scientific) and Qubit^®^ dsDNA HS assay kits (Molecular Probes Life Technologies), respectively. Quantification of OsHV-1 was carried out using quantitative PCR (qPCR). Amplification reactions were performed using the Roche LightCycler 480 Real-Time thermocycler on three technical replicates (qPHD-Montpellier GenomiX platform, Montpellier University, France). The total qPCR reaction volume was 1.5 μL, comprising 0.5 μL of DNA (40 ng/μL) and 1 μL of LightCycler 480 SYBR Green I Master mix (Roche) containing 0.5 μM PCR primers (Eurogenetec SA). The primers used were virus-specific and targeted the region of the OsHV-1 genome predicted to encode a catalytic subunit of DNA polymerase (ORF100, AY509253): Forward: 5’-ATTGATGATGTGGATAATCTGTG-3’ and Reverse: 5’- GGTAAATACCATTGGTCTTGTTCC-3’ (Webb et al., 2007). The following program was applied: enzyme activation at 95° C for 10 min followed by 40 cycles of denaturation (95°C, 10 s), annealing (60°C, 20 s) and elongation (72°C, 25 s). To check the specificity of the amplification, a subsequent melting step was applied. Twenty-one samples with a viral load greater than or equal to 10^5^ genomic units (GU)/ng of DNA were selected for sequencing (5 from Br, 7 from MO and 9 from Th). DNA-Seq library preparation and sequencing were performed by the Genome Quebec Company (Genome Quebec Innovation Center, McGill University, Montreal, Canada) using the Shotgun PCR-free library preparation kit (Lucigen) and the NovaSeq™ 6000 Sequencing system (Illumina^®^) (paired ends, 150 bp).

### Raw read QC and selection of viral reads

Read quality was assessed using FastQC v0.11.8 (Andrews, 2010), and sequence adapters were removed using trimmomatic 0.39 (Bolger et al., 2014). Reads shorter than 50 bp were discarded, and bases at the start and the end of a read were trimmed when their PHRED quality score was below 30. Additionally, reads were clipped if the average quality within a 4-bp sliding window fell below 15. PCR duplicates were removed using Picard v2.22.4 (“Picard toolkit,” 2019 http://broadinstitute.github.io/picard/). Then, viral reads were extracted using KrakenUniq 0.5.8 (Breitwieser et al., 2018) and Seqkit v0.12.0 (Shen et al., 2016) with the seq --name --only-id and grep --pattern-file. To avoid any host contamination, extracted reads were aligned to a *C. gigas* reference genome (assembly version V9 (Zhang et al., 2012)) using Bowtie2 4.8.2 (Langmead and Salzberg, 2012), and non-aligned viral reads were kept for subsequent analyses.

### Rarefaction analysis

In order to verify that OsHV-1 sequencing depth was sufficient to accurately characterise whole viral genomic diversity, we performed a rarefaction analysis for each of the 21 libraries. To this end, we performed an iterative variant calling analysis by subsampling and mapping the reads on the OsHV-1 µVar A genome (<u>GenBank: KY242785</u>).

In order to accelerate analyses, a nonlinear subsampling incrementation was performed. We started by subsampling 1000 reads until reaching 10% of the total number of reads of the library analysed. Then we subsampled 4000, 11000, 21000 and 41000 reads until reaching 30%, 50%, 80% and 100% of the total number of reads of the library respectively. After each step of subsampling/variant calling, variants were compared to the previously identified ones and the new ones were classified in SNPs and InDels in order to increment rarefaction curves.

### *De novo* assembly of non-redundant OsHV-1 genomes

Viral reads were used for *de novo* assembly using SPAdes with the --meta and --only-assembler options (Nurk et al., 2017). Assembled scaffolds were aligned to the OsHV-1 µVar A genome (GenBank no. KY242785.1) using BLAST 2.9.0 (Altschul et al., 1990) with an e-value of 0.00001. Scaffolds were then extended using SSPACE v3.0 (Boetzer et al., 2011). Considering that the OsHV-1 genome is composed of a combination of unique (U) and repeated (R) regions (TRL–UL–IRL–X–IRS–US–TRS-X, (Davison et al., 2005)) and that the latter have a size varying from 160 to 9777 bp, most of these repeat sequences could not be resolved using a kmer approach.

Scaffolds were grouped according to their size using Seqkit v0.12.0 (Shen et al., 2016). The first group of scaffolds had a size between 2 kbp and 4 kbp and made it possible to select the US region. The second group was composed of sequences with a size between 4 kbp and 20 kbp, which made it possible to select the scaffold constituting IRL-X-IRS. Finally, the last group contained sequences greater than 20 kbp, to select the UL region. We used these three groups of scaffolds to construct non-redundant genomes (NR-genomes) that contain only one copy of each repeated region (UL-IRL-X-IRS-US) (Morse et al., 2017). Because the repeat regions are inverted complements of each other, several adjustments were made to avoid the assembly of sequences in the wrong orientation. Using this strategy, 21 OsHV-1 NR-genomes were assembled from the 21 individual oyster samples.

### Whole-genome comparisons and phylogenetic analyses

The 21 NR-genomes were then aligned with the 6 NR-genomes derived from previously published OsHV-1 genomes (AY509253, KU096999, KP412538, KY242785, KY271630, MG561751, MF509813) using MAFFT v7.475 with default parameters (Katoh et al., 2002). A maximum-likelihood phylogenetic inference was conducted on the 27 NR-genomes using PhyML v3.1 (Guindon et al., 2009) with 100 bootstrap replications. We used a Hasegawa-Kishino-Yano nucleotide substitution model with invariant sites and gamma distributed categories of rate variation (HKY+G+I), which was considered as the most appropriate model by jModelTest v2.1.10 (Darriba et al., 2012) based on Akaike information criterion. Similarly, in order to get a finer picture of phylogenetic relationships among samples, the 21 NR-genomes were aligned with OsHV-1 µVar A (KY242785) genome only and a maximum-likelihood tree was obtained using an HKY+G+I substitution model. Results were locally visualized using the R package ggTree v2.0.4 (Yu et al., 2016) and FigTree v1.4.4 (available at http://tree.bio.ed.ac.uk/software/figtree/).

### Genomic variability within and between farming areas

An average non-redundant genome (A-genome) of the 21 newly generated NR-genomes was built using the “cons” argument from the EMBOSS suite v6.6.0.0 with the default settings (Rice et al., 2000). The NR-genomes were compared to the A-genome using MUMer4 4.0.0beta2 (Marcais et al., 2018) with the option “nucmer -c 100 -l 15 –f” to identify all the variable positions between the 21 NR-genomes.

To assess the genomic variability between the farming areas, both minor and major variations of each sequencing library (21) were called on the A-genome previously generated using Freebayes v1.3.2-dirty (Garrison and Marth, 2012), with the following settings: --use-mapping-quality, --min-repeat-entropy 1, --haplotype-length 0, --min-alternate-count 5, -- pooled-continuous, --hwe-priors-off, --allele-balance-priors-off. The resulting variant calling outputs were normalized using BCFtools v- (Narasimhan et al., 2016), decomposed with vt v1.0.0 (Tan et al., 2015) and split with vcflib v1.0.0. Considering the very rare occurrence of multi-nucleotide polymorphisms, these latter were counted as InDels. Variable positions with a frequency of >50% (major variations) were compared to those obtained with MUMer4 for comparative genomics and all were validated. They were subsequently subtracted from the variant calling files to retain only minor variations (with a frequency of <50%). A unique identifier composed of the position and sequence information (e.g. 128737_A>G) of each minor variation was created. Set analyses of nucleotide variations were performed with matrix approach using R (R Core Team, 2018) operation and visualized with the Venn Diagram package (Chen and Boutros, 2011), UpSetR v1.4.0 (Lex et al., 2014) or pheatmap v1.0.12.

All downstream analyses (tables, graphs, plot creation and edition) were performed in R 3.6 (R Core Team, 2018) on R studio IDE (R-Studio-Team, 2015) with an extensive use of Dplyr v1.0.0 (Wickham et al., 2018), and ggplot2 (Wickham, 2016) from the tidyverse v1.3.0 package (Wickham et al., 2019). Genome visualization was carried out using gggenes v0.4.0. For statistical analyses, samples were treated as three groups, representing the different farming areas (Br, MO and Th. Kruskal-Wallis and Dunn’s multiple comparison tests were used locally (GraphPad Prism 8.4.2) to compare the number of SNPs or InDels.

### Genomic annotations and variations effect predictions

Genomic annotation was performed using five gene prediction softwares : Prodigal (Hyatt et al., 2010), FragGeneScan (Rho *et al*., 2010 ; version 1.31), GeneMark with heuristic models ((Zhu et al., 2010) ; version 3.25) GeneMarkS for virus ((Besemer et al., 2001); version 4.28) and finally ORF finder implemented in Geneious Prime (version 2022.0.1). All identified ORFs were manually sorted in Geneious Prime and only those predicted by at least three gene predictors were retained for genomic annotation. VCF files of major and minor variations were used to generate two global VCF files containing the coordinates of the variations according to the A-genome. SnpEff ((Cingolani et al., 2012); version 4.1l) was then used to predict the effect of major and minor variations using a home-made database constructed with the A-genome and its annotations.

### Ancestral states reconstruction of discrete trait

To infer the geographic origin of our sampled isolates, sampling sites (Th, MO, Br) were modelled as a discrete trait for each NR-genome sequence over the genealogy by ancestral state inference using a discrete asymmetric phylogenetic diffusion model (Lemey et al., 2014) in BEAST v1.10.4 (Suchard et al., 2018). This approach estimates the probability of the internal nodes and branches being associated with a specific sampling site, based on information about sampling site states of the samples at the branch tips. A Bayesian stochastic search variable selection procedure (Lemey et al., 2009) was employed to allow for diffusion between specific location pairs to be included or excluded from the model. Based on Akaike’s information criterion ranking in jModelTest v2.1.10 (Darriba et al., 2012), analyses were performed under a GTR+I+G model of nucleotide substitution. A strict molecular clock and a constant coalescent model were used and a MCMC chain of 10,000,000 steps was run and sub-sampled every 1,000 generations. Convergence to the stationary distribution and sufficient mixing (effective sample size >200) for all parameter estimates were checked in Tracer after removing the initial 10% of the samples as burn-in. The BEAGLE library was used to increase computational speed (Ayres et al., 2012). Bayes Factors (BF) were computed in SpreaD3 (Bielejec et al., 2016) and used as a measure of support for identifying frequently invoked transition rates to explain the diffusion process. Support for a rate was considered substantial when BF > 3, strong if BF > 10, and decisive if >100 (Jeffreys, 1961).

To investigate the possible impact of recombination on the ancestral state reconstruction, the analysis was repeated on an alignment were genomic regions potentially impacted by recombination events were removed. To do so, we used Recombination Detection Program (RDP) version 4.97 (Martin et al., 2015), with default settings, to identify any recombination events between the OsHV-1 NR-genomes we sequenced. As evidence for recombination, we took events detected by at least two of seven different recombination detection methods implemented in RDP: RDP, MAXCHI, and GENECONV methods in primary scanning mode and the Bootscan, CHIMAERA, SisScan, and 3SEQ methods in secondary scanning mode, each with a Bonferroni-corrected P-value cut-off of 0.05. Two genomic regions were detected by RDP4 as potentially impacted by three recombination events and removed from the alignment (213 SNPs remaining after pairwise deletion at gapped sites, 46 after complete deletion) before running jModelTest, BEAST and SpreaD3 again.

Maximum clade credibility (MCC) trees were generated from tree output files from BEAST in TreeAnnotator v1.10.4, and annotated in the FigTree v1.4.4 graphical user interface (available at http://tree.bio.ed.ac.uk/software/figtree/).

### Data availability

The datasets generated from this study can be found in the SRA database BioProject accession number PRJNA681599 with submission ID SUB8642385. All the scripts generated in this study are freely available at https://github.com/propan2one/OshV-1-molepidemio.

Complementary information is available from the corresponding authors upon reasonable request.

## Acknowledgments

The present study was supported by the EU project VIVALDI (H2020 program, no. 678589) led by Ifremer; the CNRS and the University of Montpellier. JD was supported by grant from the University of Montpellier. This work also benefitted from support from the “Laboratoire d’excellence” (LabEx) CeMEB, through the exploratory research project HaploFit and the use of the “environmental genomics” facility (http://www.labex-cemeb.org/fr/genomique-environnementale-1). This work was also supported by the Ifremer Scientific Board, through the HemoVir and GT-oyster projects and by the “Fond Européen pour les Affaires Maritimes et la Pêche” (FEAMP, GESTINNOV project n°PFEA470020FA1000007). This study is set within the framework of the “Laboratoires d’Excellences (LABEX)” TULIP (ANR-10-LABX-41). The funders had no role in study design, data collection and interpretation, or the decision to submit the work for publication.

JCA, OK, BM, JD and JME were involved in the study conception and design. BM, BP, CP, JD, and JME were involved in the collection of samples and in the experimental work. JD, CP, MJ and JME were involved in bioinformatics and statistical analyses. JD, RG, CM, MJ and JME drafted the manuscript, and all authors revised and approved the final manuscript.

We thank the staff of the Ifremer stations at Argenton (LPI, PFOM) and La Tremblade (ASIM), Frédéric Girardin and his team (Plateforme des Mollusques Marins de La Tremblade PMMLT) and the Comité Régional Conchylicole de Méditerranée (CRCM) for technical support in the production of standard (NSI) oysters and transplantation experiments. We also thank Nicole Faury (Ifremer, ASIM) and Marc Leroy for technical assistance. We are also grateful to Eric Rivals (LIRMM) for fruitful discussions.

We declare that we have no competing interests.

