## Supplementary figures for "Genetic diversity and connectivity of the Ostreid herpesvirus 1 populations in France: a first attempt to phylogeographic inference for a marine mollusc disease"

**Figure S1. Phylogenetic relationship between the 21 de novo assembled and other OsHV-1 non-redundant genomes.** Shown is a midpoint rooted maximum-likelihood phylogenetic tree of the 27 OsHV-1 NR-genomes, i.e. 21 assembled from the sequencing data generated during this study and 6 derived from Ostreid herpesvirus 1 published genomes (black font). Numbers at branch nodes are bootstrap values obtained from 100 replicates and, indicate branch confidence (number of bootstrap replicates for that branch). The 21 isolates from this study are colored according to their geographic origin, Brest (Br) in blue, Marennes-Oléron (MO) in red, and Thau Lagoon (Th) in green.

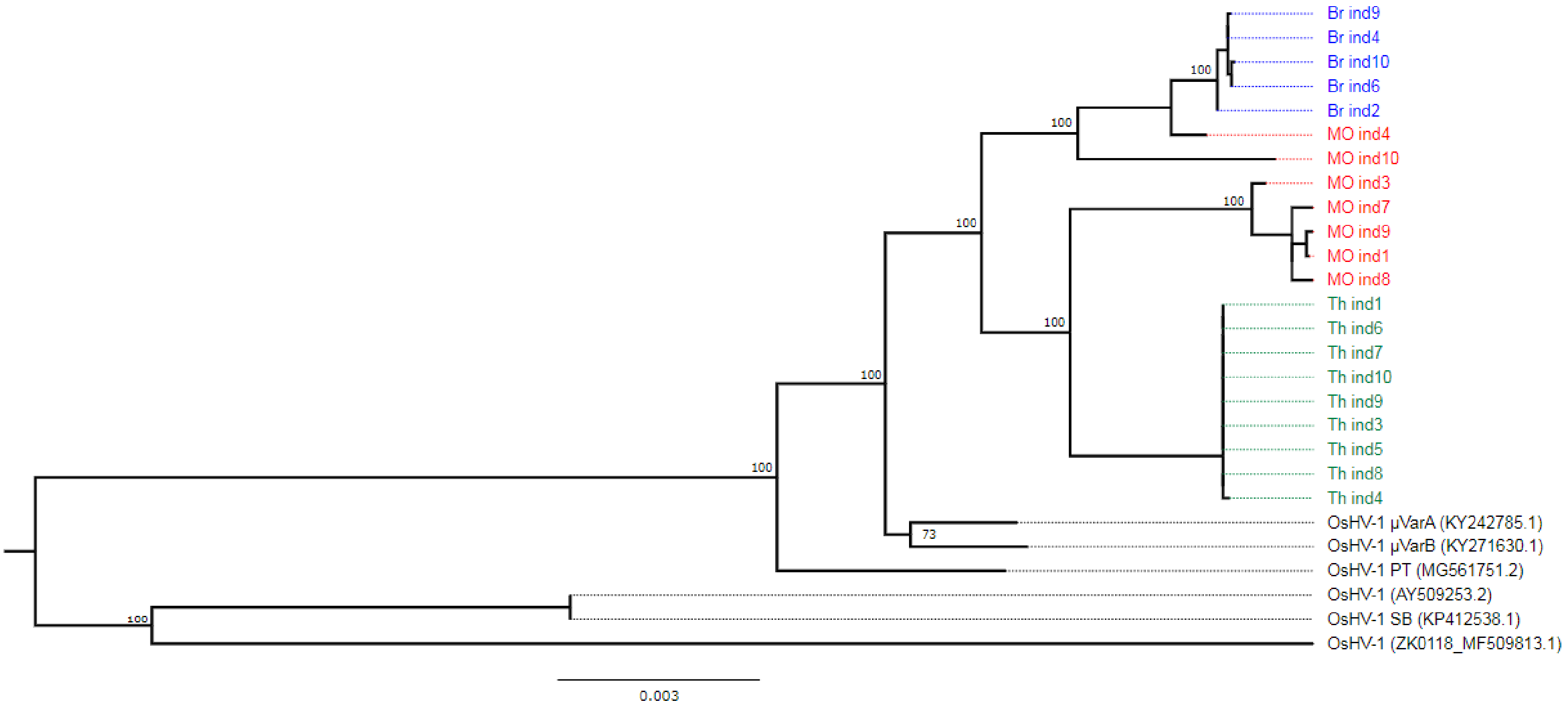

**Figure S2. Distribution of the 399 major variations across the A-genome and impact on ORF.** The upper panel represents the conventional structural regions of OsHV-1 A-genome. Grey arrow: Unique long, yellow: Inverted Repeat Long, light blue: X, dark orange: Inverted Repeat Short and purple: Unique Short. Below are represented the variations predicted to have a impact on ORFs by arrows of different colours. Dark green arrows: Unchanged ORF (not impacted by variations), orange: Frameshift variations and green: start lost. The lower panel represent the distribution and the frequency of variations across the A-NR-genome for each libraries. Colours vary according to sampling locations (Brest: blue, Marennes-Oléron: red and Thau: green) and allelic frequency (from light to dark).

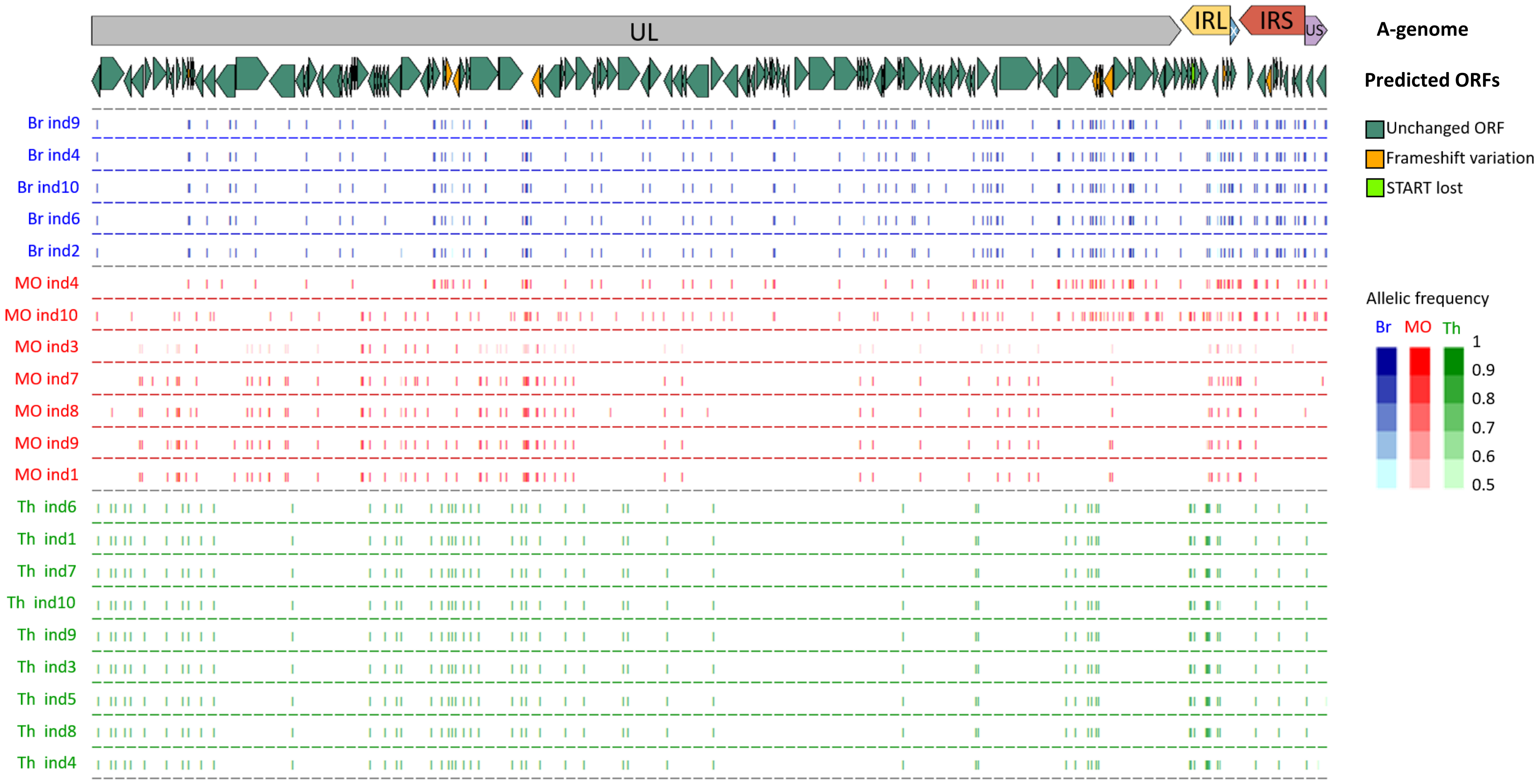

**Figure S3. Distribution of the 208 minor variations across the A-genome and impact on ORF.** The upper panel represents the conventional structural regions of OsHV-1 A-genome. Grey arrow: Unique long, yellow: Inverted Repeat Long, light blue: X, dark, dark orange: Inverted Repeat Short and purple: Unique Short. Below are represented the impact of variations on predicted ORF by arrows of different colours. Dark green arrows: Unchanged ORF (not impacted by variations), orange: Frameshift variations, burgundy red: stop gained and green: start lost. The lower panel represent the distribution and the frequency of variations across the A-NR-genome for each library. Colors vary according to sampling locations (Brest : blue, Marennes-Oléron: red and Thau: green) and allelic frequency (from light to dark).

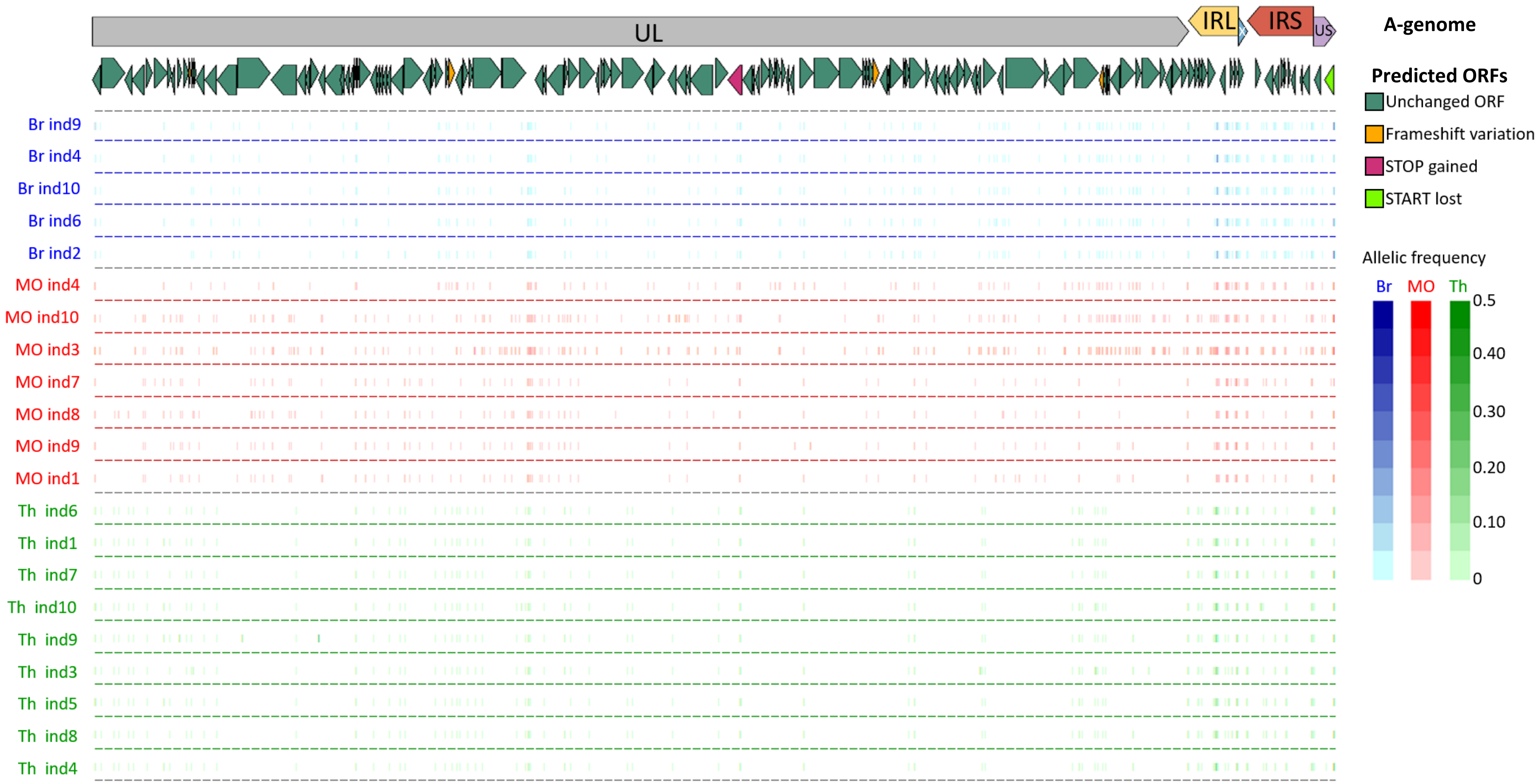

**Figure S4. Maximum clade credibility (MCC) phylogeny of NR-genomes with regions potentially impacted by recombination removed of OsVH-1 isolates and ancestral state reconstruction of the discrete locations: Brest (Br), Marennes-Oléron (MO) and Thau lagoon (Th).** Color of the branches represents the most probable location state of their descendent nodes. The color-coding is consistent with other figures: Br in blue, MO in red, and Th in green. Numbers at branch nodes indicate the posterior location probabilities. Given that we did not incorporate any timing information into our analysis (no calibration prior), the scale beneath the tree is expressed in average number of substitutions per time unit.

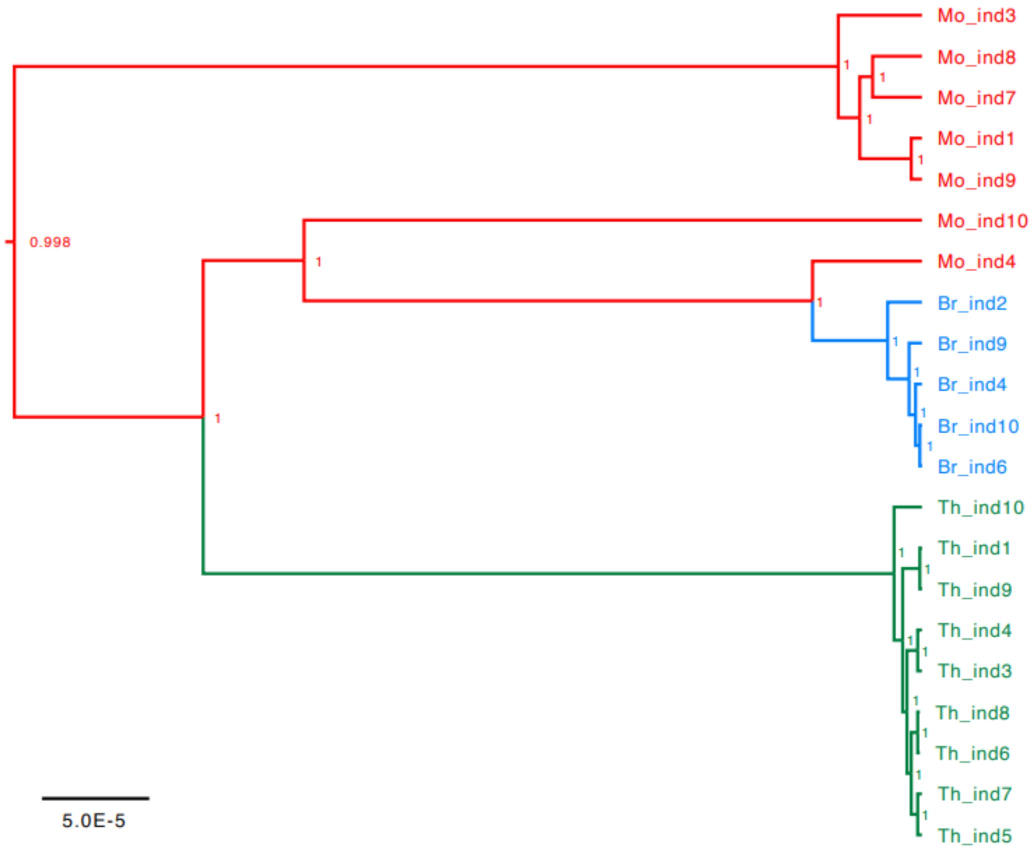
